# Proximity labelling-based identification of vascular homing peptide receptors

**DOI:** 10.1101/2025.06.14.659695

**Authors:** Rasmus Enno, Maarja Haugas, Karlis Pleiko, Kristina Põšnograjeva, Tambet Teesalu

**Affiliations:** Laboratory of Precision and Nanomedicine, Institute of Biomedicine and Translational Medicine, University of Tartu, Ravila 14b, 50411 Tartu, Estonia; Materials Research Laboratory, University of California, Santa Barbara, 93106 California, United States

**Keywords:** Tumor Homing Peptides, cell surface receptor, C-end Rule, Neuropilin-1, proximity labelling, horseradish peroxidase

## Abstract

Identifying the receptors of vascular homing peptides (VHPs) is critical for mechanistic understanding and development of peptide-guided precision therapeutics. Conventional receptor discovery methods, such as affinity chromatography, require cell disruption and often expose intracellular proteins, resulting in high background and low specificity. To overcome these limitations, we developed a proximity labelling approach that tags proteins near VHP receptors on intact live cells. Cells were incubated with VHP–horseradish peroxidase (HRP) complexes, which, upon hydrogen peroxide treatment, activate biotin-tyramide to produce short-lived radicals that covalently label nearby membrane proteins. Using the prototypic C-end Rule peptide RPARPAR and its known receptor neuropilin-1 (NRP-1), we validated this method by successful receptor tagging and mass spectrometric identification. Using RPARPAR–HRP conjugates, we achieved selective proximity labeling of membrane proteins in NRP-1–positive PPC1 cells, with a 3- to 5-fold increase in fluorescence intensity over controls by flow cytometry. Affinity purification and Western blotting identified a strong ∼130 kDa band corresponding to NRP-1 exclusively in labeled PPC1 cells. Mass spectrometry analysis revealed a ∼20-fold enrichment of NRP-1 and significant enrichment of integrins (ITGAV, ITGB1, ITGA3), ALCAM, EPHA2, CD109, and PLXNB2 in RPARPAR-labeled samples compared to controls.

This approach could be broadly used for molecular mapping of the homing peptide interactome and its spatial proximity in live cells, streamlining the discovery of homing peptide receptors and their associated partners.

## Introduction

Vascular endothelial cells (VEC) form the inner lining of blood vessels and play a central role in maintaining vascular homeostasis, regulating vascular permeability, mediating immune cell trafficking, and supporting tissue-specific signalling^7,8^. The endothelium is highly dynamic and heterogeneous, exhibiting organ-specific molecular profiles and undergoing phenotypic remodelling in response to physiological or pathological stimuli such as inflammation and tumorigenesis^6,9,10^. In cancer, angiogenesis leads to the formation of structurally and functionally abnormal blood vessels that support tumor progression, often marked by elevated levels of Vascular Endothelial Growth Factor A (VEGF-A) and its receptor VEGFR-2, as well as increased expression of NRP-1, a VEGF co-receptor that enhances pro-angiogenic signalling in tumor endothelium^11–13^. These aberrations reflect organ- and disease-specific changes in endothelial gene expression and protein composition, forming unique molecular signatures known as vascular ZIP codes, which can be leveraged for selective targeting using blood-circulating ligands^7,14–16^.

In vivo phage display is a powerful method for identifying VHPs, short ligands that navigate tissue-specific molecular ZIP codes on endothelial cells^2,17^. Libraries of phage-displayed peptides are injected systemically, and those accumulating in specific organs are recovered and sequenced to uncover homing candidates^1,18^. This strategy has yielded several notable peptides. The prototypic iRGD (CRGDKGPDC) binds αv integrins and NRP-1, enhancing tumor penetration^19^; the RPARPAR peptide utilizes the C-end Rule motif to target NRP-1 and promote vascular permeability^14^; and the CSG peptide homes to laminin–nidogen complexes in tumor extracellular matrix^20^. More recently, a peptide selectively targeting ischemic cardiac tissue was discovered using in vivo phage display combined with next-generation sequencing, demonstrating the technique’s expanding potential in precision targeting^21^. Despite these successes, phage display alone does not identify peptide receptors; deconvolution typically requires techniques like affinity purification or proximity labelling, and reliably linking peptides to their receptors remains a key bottleneck in VHP development^1,2^.

Identifying receptors for vascular homing peptides remains technically challenging due to the complex nature of membrane protein interactions and the limitations of conventional methods^3– 5^. Affinity-based techniques, such as protein pull-down from lysates, often suffer from high background caused by abundant intracellular proteins that are not normally accessible to circulating ligands^3,4^. Additionally, many receptors, such as integrins, require specific detergent conditions to preserve their native conformation during extraction^4,22^. Harsh lysis conditions may disrupt labile receptor–ligand interactions or displace co-receptors, complicating data interpretation^3,22^. To overcome these issues, various proximity labelling strategies have been developed to enable receptor identification in a more native context^5,23–25^. Among these, APEX2 and TurboID are widely used for intracellular interactome mapping, however, both enzymes are difficult to adapt for use at the extracellular membrane^5,23^. While a few studies have explored antibody-conjugated APEX2 for imaging applications, including EM labelling, these implementations remain rare and are not typically applied for extracellular proteomics due to folding and functional constraints^26,27^. TurboID, by contrast, has not been reliably used as a conjugate outside of genetically encoded contexts, as it requires intracellular expression to remain functional and has not been demonstrated to work in antibody-or ligand-conjugated forms^4,5^. Its constitutive activity and reliance on cellular cofactors such as ATP further complicate its use in spatially confined extracellular labelling^28^. PhoxID, a light-activated method, provides high spatial resolution but requires direct conjugation of a bulky photocatalyst to the ligand^24^. For peptides, which typically bind with low affinity in monovalent formats, this conjugation can reduce receptor binding and lower labelling efficiency, limiting the method’s applicability in peptide receptor discovery^29^.

In contrast, HRP-based proximity labelling offers several key advantages that make it particularly suitable for extracellular receptor discovery. HRP remains inactive in the reducing intracellular environment, inherently restricting its catalytic activity to the cell surface^30^. Upon addition of hydrogen peroxide, HRP catalyzes the oxidation of biotin-tyramide, generating short-lived phenoxyl radicals that covalently react with nearby tyrosine residues on adjacent membrane proteins^31^. This biotinylation occurs in close proximity to the HRP–ligand complex, resulting in spatially confined labelling without compromising membrane integrity^4,5,31^. The timed addition of hydrogen peroxide and biotin-tyramide allows for precise temporal control of the reaction, minimizing background and enhancing specificity^4,25^. This strategy has been successfully applied to map receptor microenvironments using antibody–HRP fusions and to identify aptamer targets through selective biotinylation^32,33^. The SPPLAT method (selective proteomic proximity labelling assay using tyramide), developed by Rees et al., demonstrated that HRP-conjugated ligands can effectively label neighbouring proteins for downstream enrichment and identification by mass spectrometry^25^. Together, these features make HRP-based proximity labelling a versatile and powerful tool for mapping membrane protein interactions under near-native conditions.

Among the vascular homing peptides identified by in vivo phage display, a distinct class known as C-end Rule (CendR) peptides has shown exceptional promise for tumor targeting and tissue penetration^14,19,34,35^. The most clinically advanced example is iRGD, which selectively accumulates in tumors and has progressed through Phase 1B/2A clinical trials to enhance intratumoral drug delivery^34^. iRGD initially binds to αv integrins and is then proteolytically processed to expose the CendR motif, enabling subsequent interaction with neuropilin-1 NRP-1 and promoting deep tissue penetration^14,19^.

Another well-characterized CendR peptide is RPARPAR, a linear hexapeptide containing a canonical C-terminal Arg–Pro–Arg motif that binds directly and specifically to NRP-1^14,36–38^. Structural studies indicate that the terminal arginine docks into the b1 domain of NRP-1, forming a salt bridge with Asp320 and stabilizing the interaction through additional electrostatic and hydrogen-bond contacts^39,40^. Upon binding, RPARPAR engages NRP-1 and triggers its recruitment into cholesterol-dependent membrane microdomains, initiating a non-canonical, receptor-directed endocytic process that is independent of clathrin, caveolin, and dynamin^14,19,41^. This pathway shares morphological features with macropinocytosis and traffics internalized cargo through the endolysosomal system, as shown for related CendR peptides^41,42^. Notably, a fraction of internalized vesicles undergoes transcytosis, allowing the peptide—and any attached cargo—to cross endothelial barriers and accumulate in the underlying tissue^41,43^. Because RPARPAR exploits this active transport mechanism while maintaining high binding specificity for NRP-1, it enables efficient and uniform delivery of conjugated or co-administered agents such as drugs, imaging probes, and nanocarriers^37,42,43^. Its robust tumor selectivity, predictable pharmacokinetics, and defined receptor interaction have established RPARPAR as a benchmark ligand for studying NRP-1–mediated vascular transport.

Building upon the unique trafficking properties of CendR peptides and the surface-specific labelling capabilities of HRP, we sought to develop a strategy for receptor identification that preserves membrane integrity and leverages native ligand–receptor interactions. Using RPARPAR as a model peptide and HRP as the enzymatic labelling moiety, we established an in vitro proximity labelling workflow capable of selectively tagging membrane proteins in live cells. To validate this approach, we applied mass spectrometry to identify biotinylated proteins. The following sections describe the experimental design, optimization steps, and validation of the HRP-mediated receptor labelling strategy.

## Results

The vascular homing peptide RPARPAR, containing the canonical CendR motif (R/KXXR/K), was synthesized with an N-terminal biotin to enable conjugation to streptavidin–horseradish peroxidase (SA-HRP), generating the proximity labelling conjugate RPAR-HRP. As a negative control, biotin without a functionalized peptide in conjunction to streptavidin–HRP (biot-HRP) was used (Fig 1A). To assess specificity, PPC1 prostate cancer cells (NRP-1 positive) and M21 melanoma cells (NRP-1 negative) were chosen as model systems.

**Figure 1.**
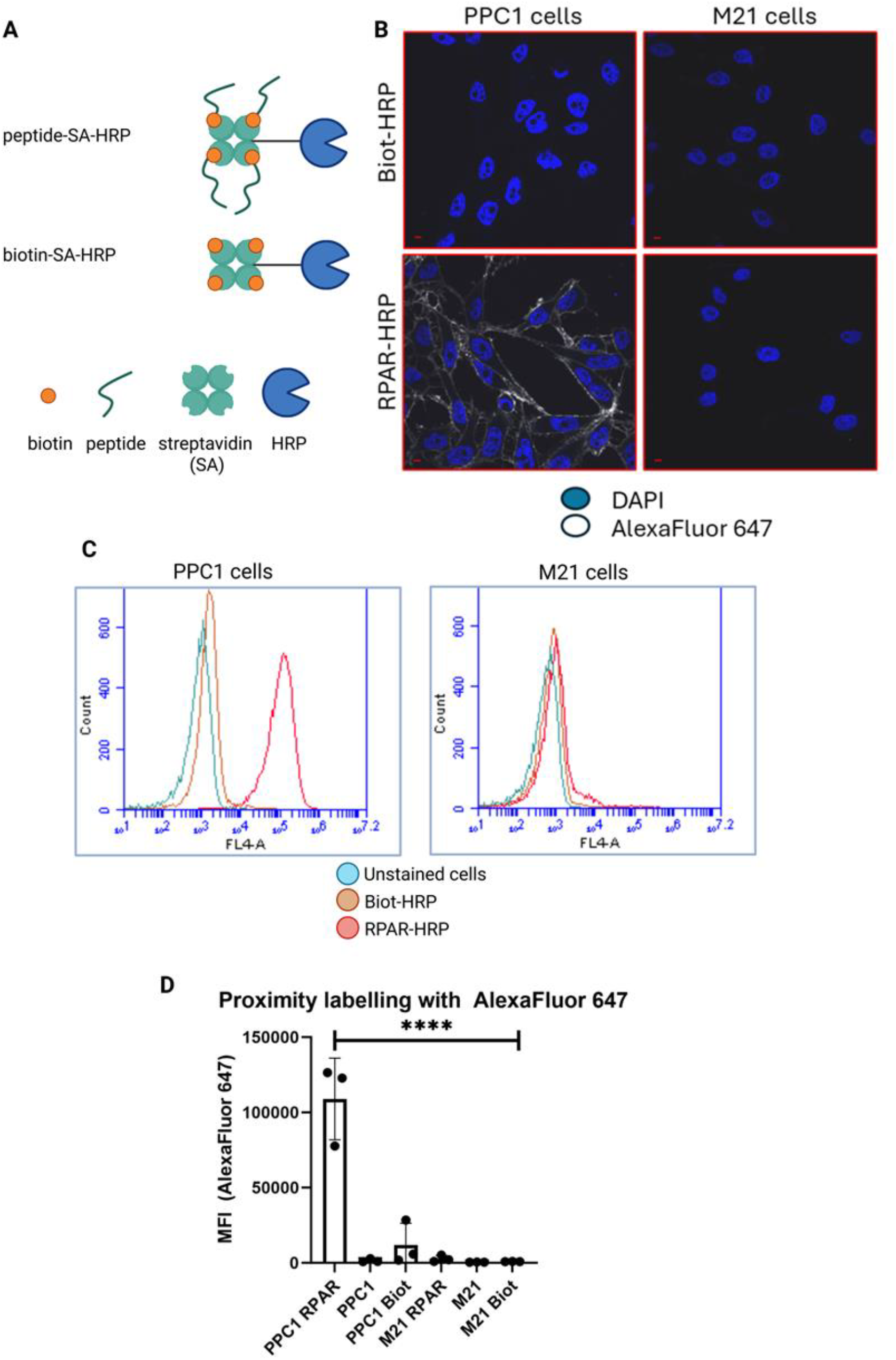
HRP-dependent proximity labelling using peptide–SA–HRP conjugates in PPC1 and M21 cells. **A**. Schematic representation of peptide–SA–HRP conjugates. Created with BioRender.com. **B**. Confocal microscopy images of PPC1 and M21 cells labelled via HRP-dependent proximity ligation using AlexaFluor™ 647–tyramide. Cells were incubated with RPAR–HRP or biotin–HRP, followed by proximity labelling with AlexaFluor 647–tyramide (white). Nuclei were counterstained with DAPI (blue). White signal indicates successful cell surface labelling via the HRP–tyramide reaction. Scale bar: 15 µm. Images were acquired using an Olympus FV1200MPE confocal microscope with a 60× oil-immersion objective. Representative results from three independent experiments (n = 3; Supplementary figure 1) **C**. Flow cytometry analysis of PPC1 and M21 cells labelled with AlexaFluor™ 647–tyramide. Cells were incubated with RPAR–HRP (red), biotin–HRP (brown), or left untreated (light blue), followed by proximity labelling. Histograms display AlexaFluor 647 fluorescence intensity (x-axis) versus cell count (y-axis). Data are representative of three independent experiments (n=3; Supplementary figure 2). **D**. Quantification of AlexaFluor 647 signal by mean fluorescence intensity (MFI). Bar graph shows mean FL-A values for PPC1 and M21 cells treated with RPAR–HRP, biotin–HRP, or left untreated. Data represent mean ± standard deviation (SD) from three independent experiments (n = 3). A statistically significant increase in fluorescence was observed in PPC1 cells treated with RPAR–HRP. One-way ANOVA followed by Tukey’s HSD post hoc test was used for multiple comparisons. Statistical significance was defined as *p* < 0.05; **** indicates *p* < 0.001 versus untreated PPC1 cells.

To evaluate the efficiency and selectivity of proximity labelling, cells were treated with RPAR-HRP or biot-HRP, followed by a 2-minute incubation with AlexaFluor 647-tyramide and hydrogen peroxide. Confocal microscopy revealed a strong AlexaFluor 647 signal localized to the plasma membrane in PPC1 cells treated with RPAR-HRP, whereas M21 cells or cells treated with biot-HRP exhibited only background levels of fluorescence (Fig. 1B). This demonstrates that the enzymatic activity of HRP and the targeting selectivity of RPARPAR are preserved after conjugation, and that the conjugate can be used to selectively label NRP-1–positive cells.

Flow cytometry analysis further quantified this labelling effect. In PPC1 cells, treatment with RPAR-HRP induced a marked increase in fluorescence intensity compared to biot-HRP and unstained controls. In contrast, M21 cells exhibited no discernible shift in fluorescence intensity under any condition, confirming that the signal is NRP-1 dependent (Fig 1B, 1C). Moreover, the statistical analysis showed that PPC1 cells treated with RPAR-HRP achieved statistically significantly higher mean fluorescence intensity (MEAN FL-A) than all the other samples (Fig 1D). These results demonstrate that the enzymatic activity of HRP and the targeting selectivity of RPARPAR are preserved after conjugation, and that the conjugate can be used to selectively label NRP-1–positive cells. Additionally, the 2-minute proximity ligation reaction time is sufficient to achieve labelling with AlexaFluor 647-tyramide.

### Biotin-Tyramide Enables Labelling for Affinity Purification

To facilitate downstream enrichment and proteomic identification, we replaced the fluorescent tyramide with biotin-tyramide. Following incubation with RPAR-HRP and Biot-HRP and the proximity ligation reaction, cells were stained with streptavidin-DyLight 550 and analysed by flow cytometry. Again, only RPAR-HRP-treated PPC1 cells showed significant labelling, with minimal signal in M21 or biot-HRP-treated cells (Fig. 2A, 2B). These findings validate that the proximity labelling system is compatible with both fluorescent and affinity-tagged tyramide derivatives.

**Figure 2.**
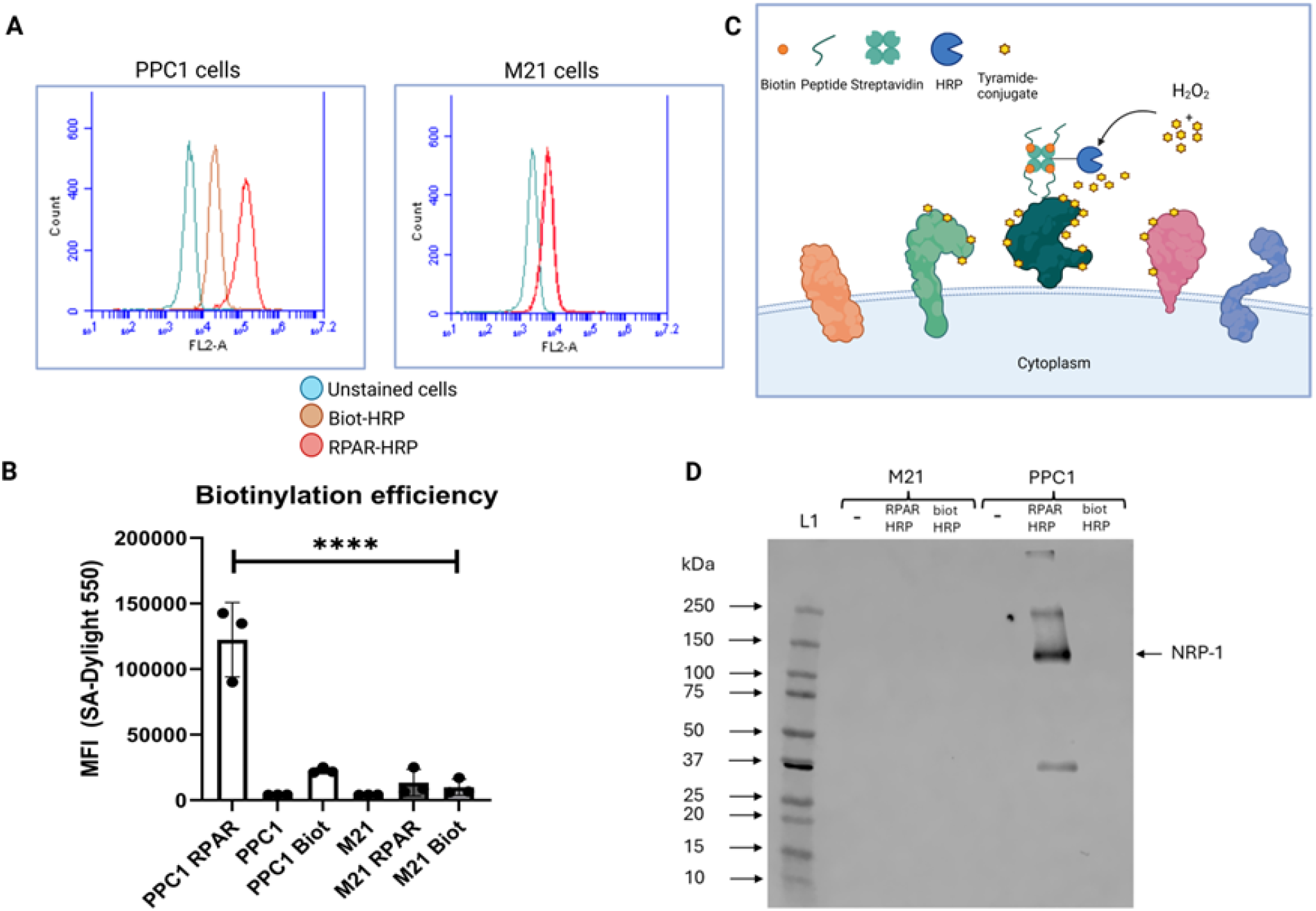
HRP-dependent proximity labelling with biotin–tyramide in PPC1 and M21 cells. **A**. Flow cytometry analysis of PPC1 and M21 cells labelled via HRP-mediated proximity biotinylation. Cells were incubated with RPAR–SA–HRP (red), biotin–SA–HRP (brown), or left untreated (light blue). Proximity labelling was performed using biotin–tyramide, followed by staining with streptavidin–DyLight™ 550 to detect biotinylated surface proteins. Histograms show streptavidin–DyLight 550 fluorescence intensity (x-axis) and cell count (y-axis). Data are representative of three independent experiments (n = 3, supplementary figure 3). **B**. Quantification of proximity labelling with biotin–tyramide. Bar graph shows the Mean FL-A of PPC1 and M21 cells after treatment with RPAR–SA–HRP, biotin–SA–HRP, or no treatment. Data represent the mean ± standard deviation from three independent experiments. Statistical analysis was performed using one-way ANOVA followed by Tukey’s HSD post hoc test. A p-value < 0.05 was considered statistically significant. **** indicates p < 0.001 compared to untreated PPC1 cells. **C**. Schematic overview of HRP-dependent proximity labelling with biotin–tyramide. Upon binding of the peptide–HRP complex to the cell surface receptor, the addition of biotin–tyramide and hydrogen peroxide initiates a catalytic reaction. HRP catalyzes the oxidation of the tyramide moiety, generating a short-lived radical that covalently labels nearby proteins via tyrosine residues. This reaction enables spatially restricted labelling of membrane-associated proteins in proximity to the targeting peptide. **D**. Western blot analysis of biotinylated proteins following HRP-dependent proximity labelling. PPC1 and M21 cells were incubated with RPAR–HRP, biotin–HRP, or left untreated, followed by proximity labelling with biotin–tyramide. Biotinylated proteins were enriched via streptavidin-based affinity purification, separated by SDS-PAGE, and transferred to PVDF membranes. NRP-1 detection was performed using a rabbit monoclonal anti-NRP-1 antibody. Lane L1 represents the molecular weight marker.

Biotinylated membrane proteins from PPC1 and M21 cells were isolated using magnetic Strep-Tactin beads. Eluted fractions were subjected to SDS-PAGE. These results were validated by Western blot using an anti-NRP-1 monoclonal antibody. A prominent band at ∼130 kDa, corresponding to NRP-1, was observed only in RPAR-HRP treated PPC1 samples; it was not detectable in PPC1 cells treated with biot-HRP or left untreated, nor in any M21 samples.

### Mass Spectrometry Validation

To identify the captured proteins, eluates were analysed by liquid chromatography-tandem mass spectrometry (LC-MS/MS). NRP-1 was among the top hits, confirming the specificity and functionality of the method. NRP-1 was among the top hits after statistical analysis, with a log_2_ fold change (log_2_FC) of 4.32 (∼20-fold increase). Beyond NRP-1, other proteins such as integrin alpha-V (ITGAV), integrin beta-1 (ITGB1), integrin alpha-3 (ITGA3), activated leukocyte cell adhesion molecule (ALCAM), ephrin type-A receptor 2 (EPHA2), CD109, and Plexin-B2 (PLXNB2) were significantly enriched in RPAR-HRP-treated PPC1 samples.

## Discussion

Our findings indicate that peptide-guided, HRP-dependent proximity labelling enables selective and efficient identification of membrane receptors in intact cells under near-physiological conditions. HRP to the CendR peptide RPARPAR, we developed a system that facilitates temporally controlled, spatially confined labelling of proximal proteins at the cell surface. This strategy effectively bypasses the limitations of traditional pull-down methods, which rely on cell lysis and are prone to capturing non-physiological interactions with intracellular components^3,4^. Unlike HRP, other commonly used proximity labelling enzymes such as TurboID and APEX2 are primarily optimized for intracellular applications^4,5,23^. They either require genetic encoding, rely on intracellular cofactors (e.g., ATP), or produce diffusible reactive species that can compromise spatial precision^5,23^. These features limit their utility in extracellular or ligand-guided receptor mapping workflows, particularly where precise surface localization and temporal control are essential.

The RPAR-HRP conjugate retained both peptide binding specificity and peroxidase activity following conjugation. Confocal microscopy and flow cytometry experiments confirmed that RPAR-HRP selectively binds and labels NRP-1–expressing PPC1 cells with AlexaFluor 647-tyramide, while showing minimal signal in NRP-1–negative M21 cells or in Biot-HRP incubated control treatments (Fig 1B-D). These results demonstrate that both components of the conjugate function as intended, and that proximity labelling is restricted to the surface of target cells. Notably, fluorescence localized primarily to the plasma membrane, consistent with the known inactivity of HRP in reducing intracellular environments, suggesting that labelling is confined to the cell surface and does not proceed after endocytosis.

To assess the applicability of the system for downstream affinity-based proteomic workflows, we substituted fluorescent tyramide with biotin-tyramide and first validated biotinylation efficiency by flow cytometry. Biotin signal was strongest in PPC1 cells treated with RPAR-HRP, while M21 cells and all biot-HRP controls showed minimal or background-level labelling (Fig 2A, 2B). These results confirmed that the RPAR-HRP conjugate enables selective and robust biotinylation of surface proteins in an NRP-1–dependent manner. A slight increase in nonspecific signal, particularly with streptavidin-DyLight detection, underscores the need for careful washing and appropriate controls. Following this validation, biotinylated proteins were isolated from detergent-lysed cells using Strep-Tactin XT magnetic beads for subsequent analysis.

Western blot analysis of affinity-enriched samples confirmed the isolation of NRP-1 following proximity labelling with RPAR-HRP (Fig 2D). Using an anti-NRP-1 antibody, we detected a specific ∼130 kDa band exclusively in PPC1 cells treated with the targeting conjugate, confirming the presence of NRP-1 and validating the specificity of the labelling and enrichment workflow. As expected, this signal was absent from all biot-HRP–treated and NRP-1–negative control samples. While Western blotting provides valuable protein-level confirmation, it is inherently limited by its reliance on available antibodies and cannot be used to identify unknown receptors without prior target information.

To overcome the limitations of antibody-based detection, we applied mass spectrometry to unbiasedly profile the biotinylated surface proteome. Among the proteins enriched in RPAR-HRP–treated PPC1 cells, NRP-1 was one of the top-ranking hits based on both log_2_FC (∼4.3) and FDR, confirming selective labelling and successful receptor identification (Fig 3). In addition to NRP-1, several other membrane proteins were significantly enriched. ALCAM, a known cell adhesion receptor involved in endothelial–tumor cell interactions, has previously been shown to co-localize with NRP-1 and mediate leukocyte transmigration and has been shown to play a key role in the binding and uptake of cancer-derived extracellular vesicles. PLXNB2, a receptor in semaphorin signalling, is implicated in vascular remodelling and has also been functionally associated with NRP family members. ITGB1 and ITGA3 are components of integrin heterodimers known to mediate cell–matrix interactions, and ITGB1 has been linked to NRP-1– dependent adhesion and angiogenesis.

**Figure 3.**
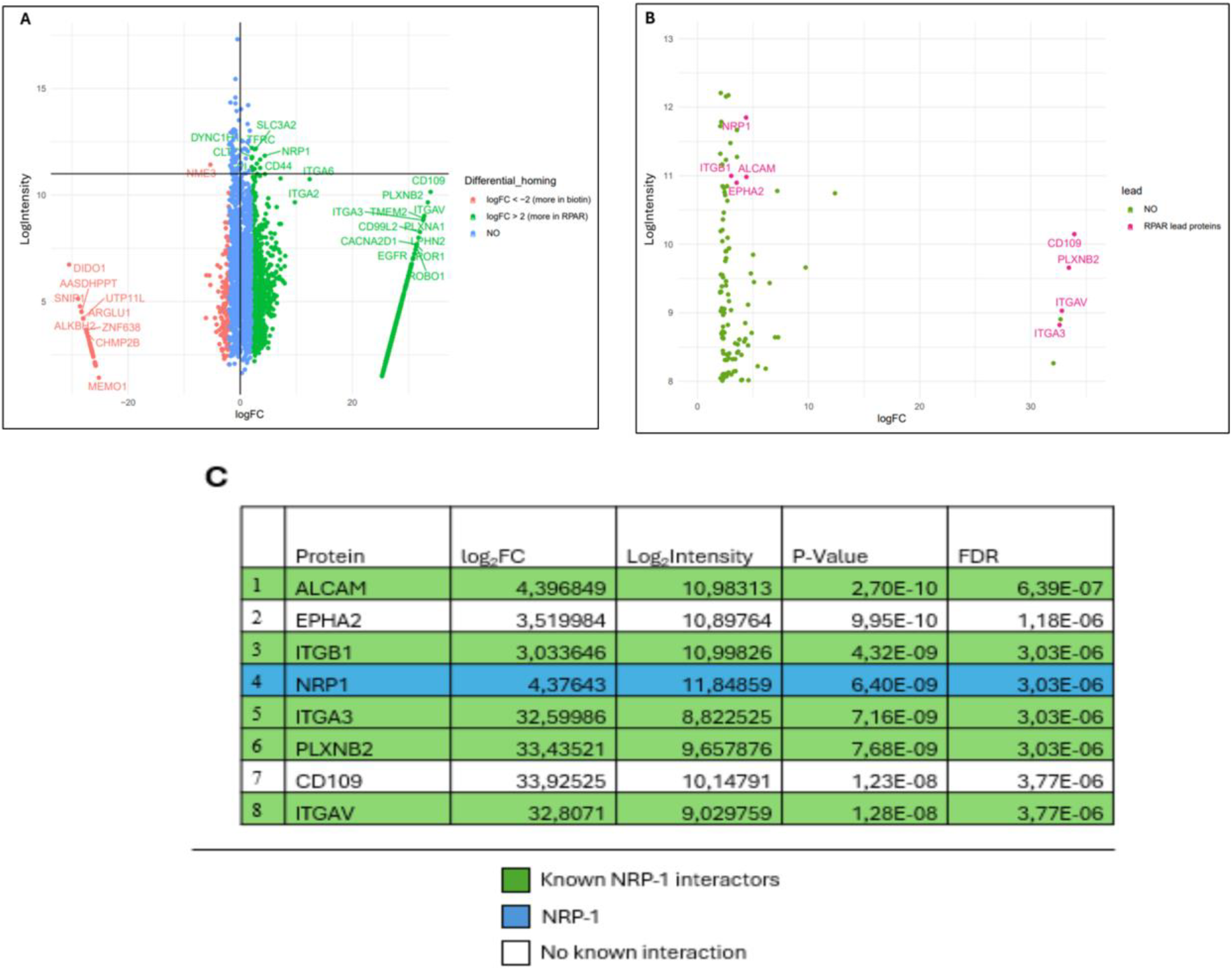
Mass spectrometry-based identification of proteins enriched by RPAR–HRP proximity labelling. **A**. Volcano plot showing the distribution of proteins detected in samples treated with RPAR– HRP (n = 4) or biotin–HRP (n = 4). Each point represents an individual protein, plotted according to log_2_FC and log_2_ intensity. Proteins significantly enriched in RPAR–HRP-treated samples (log_2_FC > 2) are highlighted in green, whereas proteins enriched in biotin–HRP-treated control samples (log_2_FC < –2) are shown in red. Proteins with no substantial difference in abundance between conditions are shown in blue. **B**. Magnified view of the upper-right region of the volcano plot, highlighting proteins highly enriched in RPAR–HRP-treated samples. Selected lead candidates are annotated in pink. **C**. Summary table of the top eight differentially abundant proteins identified in RPAR–HRP samples, ranked by false discovery rate (FDR). NRP-1 is highlighted in blue. Proteins previously reported in the literature as NRP-1 interactors are marked in green; proteins with no known interaction with NRP-1 are shown in white.

Additionally, we identified novel candidate proteins that have not been directly associated with NRP-1 but are functionally compelling. EPHA2, a receptor tyrosine kinase involved in cell repulsion, migration, and angiogenesis, shares spatial and functional overlap with NRP-1 signaling pathways. CD109, a GPI-anchored cell surface glycoprotein, is a known regulator of TGF-β signalling and is frequently overexpressed in metastatic cancers; its enrichment may reflect localization within lipid raft–associated signalling hubs. While direct interactions between NRP-1 and these proteins have not yet been demonstrated, their consistent co-enrichment suggests potential spatial proximity or complex formation at the cell surface. These findings extend the known landscape of proteins associated with RPARPAR engagement and open new avenues for dissecting NRP-1–centered signalling architectures in endothelial and tumor contexts.

Interestingly, a subset of proteins—including ITGA3, CD109, PLXNB2, and ITGAV—showed exceptionally high fold changes (log_2_FC > 30), largely due to their complete absence in control samples. This emphasizes the sensitivity of the proximity labelling approach and its capacity to capture proteins enriched specifically in the peptide-labelled microenvironment. Nonetheless, such extreme values warrant careful validation to exclude technical artifacts, such as low-abundance proteins falling below detection thresholds in controls.

The discovery that multiple proteins are co-enriched with NRP-1 supports the hypothesis that vascular homing peptides may not bind to single receptors in isolation but rather interact with dynamic receptor complexes or lipid raft–associated microdomains. This model aligns with prior reports of multivalent or co-receptor–mediated uptake mechanisms in endothelial cells and suggests that receptor identity may be context-dependent. As a result, the ability to label the peptide’s entire receptor microenvironment offers a significant advantage over traditional methods focused on individual binding partners.

While our study establishes the utility of HRP-mediated proximity labelling for receptor identification, it also reveals important limitations. The method requires prior knowledge of a peptide’s target tissue or cell line, and receptor expression may vary significantly between in vitro models and in vivo environments. Moreover, MS results show a high number of co-enriched proteins, which could be reduced with further optimization of the protocol.

Looking forward, this method can be extended to other vascular homing peptides to validate known interactions and uncover new receptor candidates. The core workflow, peptide-guided labelling, affinity enrichment, and mass spectrometry, can be adapted for different targeting motifs and detection chemistries. Of particular interest is the application of this approach in ex vivo or in vivo systems, where it could overcome the limitations of cell culture and capture receptor interactions under physiological conditions. The ability to label receptors in intact tissues would significantly advance the field of targeted drug delivery and improve our understanding of endothelial heterogeneity and peptide biodistribution.

In conclusion, our results demonstrate that peptide-directed, HRP-based proximity labelling, coupled with affinity purification and mass spectrometry, constitutes a powerful, modular strategy for vascular homing peptide receptor identification. Its application to additional peptides and biological systems will further refine our understanding of vascular targeting mechanisms and support the development of precision delivery strategies in nanomedicine.

## Methods

### Cell Lines

Primary Prostate Carcinoma-1 (PPC1) and UCLA-SO-M21 (M21) cell lines were cultured in DMEM (Lonza, BE12-604F) supplemented with 10% fetal bovine serum (FBS, Caprocorn, FBS-11A), and 1% penicillin–streptomycin at 37 °C in a humidified incubator with 5% CO_2_.

### Preparation of SA-HRP Conjugates

Biotin-ahx-RPARPAR-OH (RPAR; TAG Copenhagen) and biotin (Sigma-Aldrich, B4501) were each dissolved in Dulbecco’s phosphate-buffered saline (DPBS; Corning, 21-031-CV) to prepare 2 mM stock solutions. Streptavidin–horseradish peroxidase (SA-HRP; Sigma-Aldrich, S5512) was reconstituted in 500 µL of DPBS, and the concentration was adjusted to ≥0.5 mg/mL, as measured using a NanoDrop™ 2000c spectrophotometer (Thermo Scientific, ND-2000) with the *Protein A280* setting. This corresponds to an approximate molarity of 4.8 µM.

For the conjugation reaction, a 10-fold molar excess of either RPAR or biotin was added to SA-HRP. The mixture was incubated for 1 hour at room temperature (RT) on an end-over-end rotator. To remove unbound peptide or biotin, the reaction mixture was loaded onto a pre-wetted Amicon® Ultra centrifugal filter unit with a 3 kDa molecular weight cut-off (MWCO; Sigma-Aldrich, UFC5003). The sample was washed three times by filling the unit with DPBS and centrifuging at 14,000 × g for 10 minutes at RT, discarding the flow-through after each spin.

Following the final wash, the retained conjugate was resuspended in 50 µL of DPBS and transferred to a clean 1.5 mL microcentrifuge tube. The final concentration of the conjugate was quantified using the *Protein A280* setting on the spectrophotometer.

### Proximity Labelling and Microscopy Sample Preparation

Circular glass coverslips were placed into a 24-well plate, and 150,000 M21 or PPC1 cells were seeded per well. Cells were cultured overnight at 37 °C in a humidified incubator with 5% CO_2_. The following day, the culture medium was removed, and cells were incubated with 0.5 µM RPAR–HRP or biotin–HRP in DMEM for 1 hour at room temperature (RT).

After incubation, cells were washed three times with DPBS to remove unbound conjugate. Proximity labelling was performed by incubating the cells for 2 minutes at RT with AlexaFluor™ 647–tyramide (Thermo Fisher Scientific, B40958; 1:600 dilution in DPBS) and 1 mM hydrogen peroxide (H_2_O_2_; Honeywell, 008-003-00-9). The reaction was quenched by adding 100 U/mL catalase (Sigma-Aldrich, C9322) in DPBS, followed by three washes with catalase-containing DPBS to ensure complete termination of the reaction.

Cells were then fixed with 4% paraformaldehyde (PFA) in PBS for 10 minutes at RT and washed three times with DPBS to remove residual PFA. Nuclei were stained with 1 µg/mL DAPI in DPBS for 3 minutes, followed by two additional DPBS washes. Coverslips were mounted onto glass slides using Fluoromount-G™ mounting medium (Thermo Fisher Scientific, 00-4958-02), protected from light, and allowed to dry overnight at RT. Samples were imaged the following day using confocal microscopy.

### Flow Cytometry-Based Proximity Labelling

PPC1 and M21 cells were cultured in T75 flasks until ∼90% confluency. Cells were washed with DPBS (Corning, 21-031-CV) and detached using Cellstripper® (Corning, 25-056-CI), a non-enzymatic dissociation solution used to preserve cell surface receptors. A total of 900,000 cells per condition were transferred into wells of a 96-well V-bottom plate. After centrifugation (5 minutes, 300 × g, RT), the medium was aspirated and cells were resuspended in 0.5 µM RPAR–HRP or biotin–HRP in DMEM (Lonza, BE12-604F). Cells were incubated for 1 hour at room temperature. For autofluorescence controls, additional wells were incubated with DMEM alone.

Following incubation, cells were washed three times with DPBS (centrifugation at 300 × g, 5 minutes, RT). For fluorophore-based detection, the cell pellet was resuspended in AlexaFluor™ 647–tyramide (Thermo Fisher Scientific, B40958; 1:600 in DPBS) containing 1 mM H_2_O_2_ (Honeywell, 008-003-00-9), and incubated for 2 minutes at room temperature. For biotin-based detection, the same procedure was performed using 500 µM biotin–tyramide (Iris Biotech, LS-3500) instead of the fluorophore-labelled tyramide. In both cases, the reaction was quenched by adding 100 U/mL catalase (Sigma-Aldrich, C9322) in DPBS, followed by three additional washes using the same catalase-containing buffer to ensure complete inactivation of residual peroxide. Cells were then fixed in 4% paraformaldehyde (PFA) in PBS for 10 minutes at room temperature. To minimize cell loss during subsequent washes, samples were transferred from the 96-well plate to 1.5 mL microcentrifuge tubes. Excess PFA was removed by washing cells three times with DPBS, as described above. For biotinylated samples, cells were further incubated with Streptavidin–DyLight™ 550 (Thermo Fisher Scientific, 84542; 1:400 dilution in DPBS) for 1 hour at room temperature, followed by three washes in DPBS to remove unbound fluorophore. Finally, cells were resuspended in Milli-Q water and analysed by flow cytometry. Gating was performed based on the negative control (DMEM-treated) samples to define the background autofluorescence.

### Strep-Tactin Affinity Purification of Biotinylated Proteins

PPC1 and M21 cells were cultured in 150 cm^2^ flasks until ∼90% confluency. The culture medium was aspirated, and cells were washed once with DPBS. After removing the wash buffer, cells were detached using the non-enzymatic dissociation reagent Cellstripper® (Corning, 25-056-CI). A total of 900,000 PPC1 or M21 cells were transferred to 1.5 mL microcentrifuge tubes and incubated in DMEM (Lonza, BE12-604F) containing 0.5 µM RPAR–HRP, biotin–HRP, or no HRP (control) for 1 hour at room temperature. The labelling reaction was carried out using biotin-tyramide as described before and was quenched by adding 100 U/mL catalase in DPBS.

Cells were lysed in lysis buffer composed of 1% n-dodecyl-β-D-maltoside (DDM; Thermo Fisher Scientific, 89902), one cOmplete™ protease inhibitor tablet (Thermo Fisher Scientific, A32965) per 2 mL, and 1× sample buffer (Invitrogen, BN2008) in DPBS. Samples were mixed thoroughly and incubated on ice for 30 minutes. Following lysis, samples were centrifuged at 16,000 × g for 30 minutes at 4 °C. The supernatant (cell lysate) was carefully collected into fresh tubes.

To isolate biotinylated proteins, 20 µL of MagStrep® Strep-Tactin® XT magnetic beads (Iba Lifesciences, 2-5090-002) were added to each lysate. Samples were incubated for 30 minutes at room temperature with rotation. The beads were then washed four times with lysis buffer, transferring them to a new 1.5 mL tube after each wash.

For elution, the magnetic beads were resuspended in 30 µL of lysis buffer containing 50 mM biotin and heated at 95 °C for 5 minutes. Beads were magnetically separated, and the eluates were collected and stored at −20 °C until further analysis.

### SDS-PAGE

Eluted samples were thawed and mixed with 10 µL of loading buffer (4× Laemmli buffer containing 1:10 2-mercaptoethanol). Samples were denatured at 95 °C for 5 minutes and loaded onto 4–20% Mini-PROTEAN® TGX™ Precast Protein Gels (Bio-Rad, 4561095). Electrophoresis was performed in 1× Tris/glycine running buffer (25 mM Tris, 192 mM glycine, 0.1% SDS) at 100 V for 90 minutes.

### Silver Staining

After electrophoresis, gels intended for silver staining were fixed overnight at room temperature in fixing solution (50% methanol, 12% acetic acid, and 0.0185% formaldehyde in Milli-Q water). Gels were washed twice with 50% ethanol for 10 minutes each and pretreated for 1 minute with 200 mg/L sodium thiosulfate pentahydrate (Na_2_S_2_O_3_·5H_2_O), followed by two washes with Milli-Q water. Staining was performed by incubating the gels for 10 minutes in a solution containing 2.5 g/L silver nitrate and 0.0275% formaldehyde. After rinsing with Milli-Q water, gels were developed using developing solution (60 g/L sodium carbonate, 1.4 mg/L Na_2_S_2_O_3_·5H_2_O, and 0.0185% formaldehyde) for 5–15 minutes. Once the desired signal intensity was reached, the reaction was stopped with stop solution (50% methanol and 12% acetic acid) for 5 minutes and gels were rinsed with Milli-Q water.

### Western Blotting

For immunodetection, proteins were transferred from SDS-PAGE gels to polyvinylidene difluoride (PVDF) membranes using the Trans-Blot® Turbo™ Transfer System (Bio-Rad, 1704150) with 0.2 µm PVDF transfer packs (Bio-Rad, 1704156). The preprogrammed “Mixed MW (Turbo)” protocol was used. Membranes were washed once with 1× TBS-T (Tris-buffered saline with 0.1% Tween-20) and blocked for 30 minutes at room temperature in 5% non-fat milk in TBS-T. Blots were incubated overnight at 4 °C with a rabbit monoclonal anti-NRP-1 primary antibody (Abcam, ab81321; 1:1000 dilution in 3% milk in TBS-T), followed by a 1-hour incubation with an HRP-conjugated anti-rabbit secondary antibody (BioLegend, 406401; 1:5000 dilution in 3% milk in TBS-T). Chemiluminescent detection was performed using SuperSignal™ West Pico PLUS substrate, and membranes were imaged using the LI-COR® Odyssey® Fc Imager.

### Mass Spectrometry Analysis

PPC1 and M21 cells were incubated with 0.5 µM RPAR–SA-HRP or biotin–SA-HRP in 1.5 mL microcentrifuge tubes. Proximity labelling was performed as described above and quenched with 100 U/mL catalase in DPBS. Following centrifugation (5 minutes, 300 × g, RT), the cell pellets were snap-frozen in liquid nitrogen and stored at −80 °C.

Affinity purification of biotinylated proteins and subsequent mass spectrometry analysis were performed by the Biomolecular Mass Spectrometry and Proteomics Facility at the University of California, San Diego (UCSD). The Mass spectrometry results were analysed by Karlis Pleiko in University of Tartu

## Author contributions

Rasmus Enno participated in the study design planning, wrote the manuscript, made the figures, performed all experiments (besides Mass Spectrometry), conducted all statistical analyses (besides Mass Spectrometry) and performed or assisted in data analysis. Maarja Haugas and Kristina Põšnograjeva planned and conducted the Mass Spectrometry experiment. Karlis Pleiko analysed the mass spectrometry data. Tambet Teesalu supervised the study, participated in the study design planning and writing of the manuscript. All authors read and approved the manuscript.

