## Supplemental figures for "Proximity labelling-based identification of vascular homing peptide receptors"

|  | Biot-HRP | RPAR-HRP |
| --- | --- | --- |
| PPC1 | 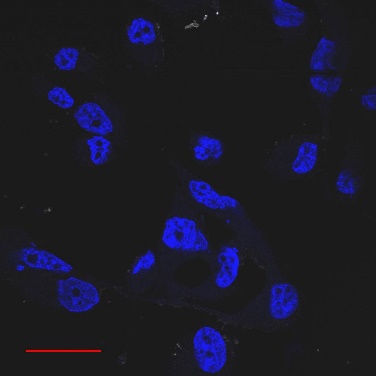 | 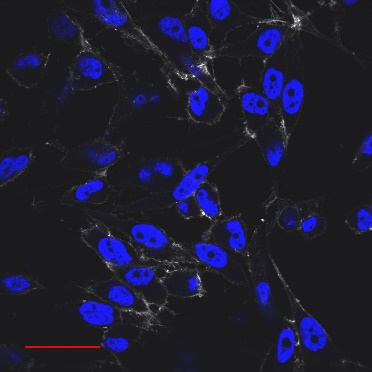 |
| PPC1 | 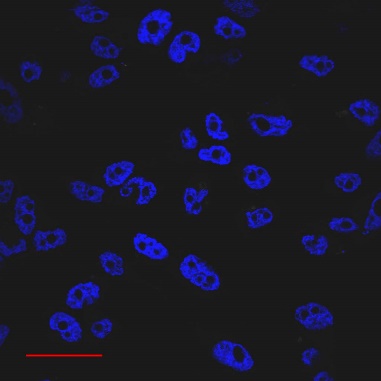 | 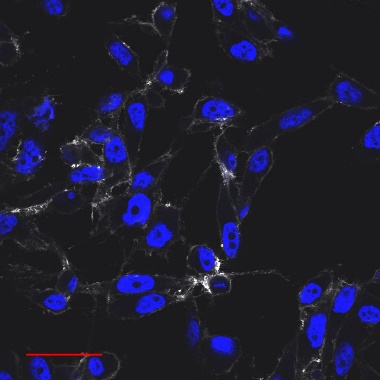 |
| M21 | 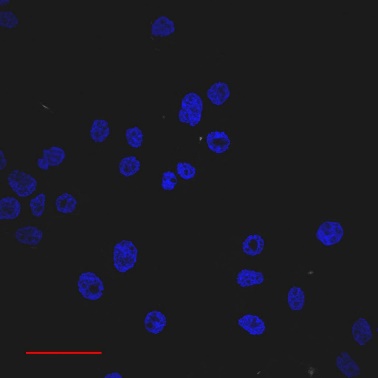 | 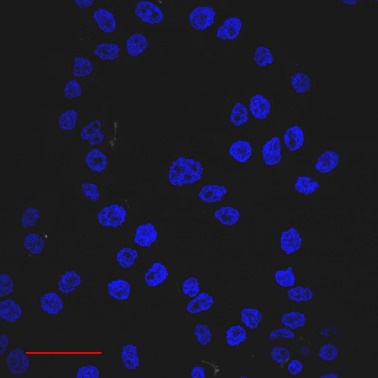 |
| M21 | 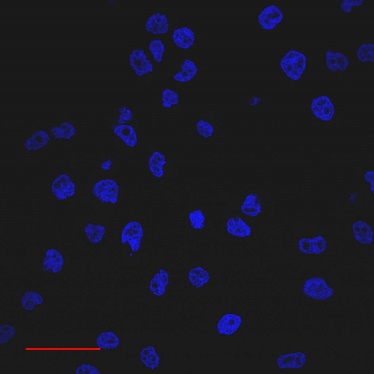 | 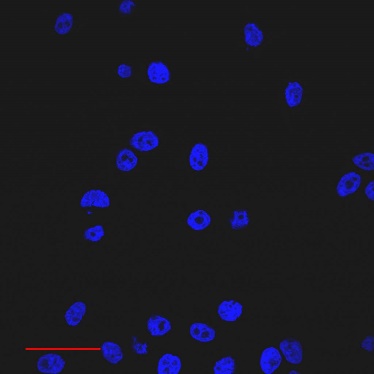 |

**Supplementary Figure 1.** Additional confocal microscopy images of PPC1 and M21 cells labelled via HRP-dependent proximity ligation using AlexaFluor™ 647–tyramide. Cells were treated and imaged as described in the main figure. Scale bar: 15 µm.

| PPC1 | M21 |
| --- | --- |
| 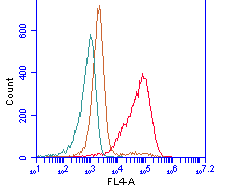 | 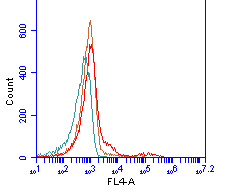 |
| 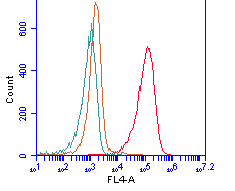 | 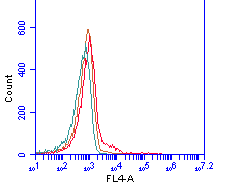 |

**Supplementary Figure 2.** Additional flow cytometry histograms of PPC1 and M21 cells labelled with AlexaFluor™ 647–tyramide following proximity labelling. Cells were treated as described in the main figure. (n=3)

| PPC1 | M21 |
| --- | --- |
| 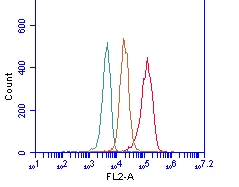 | 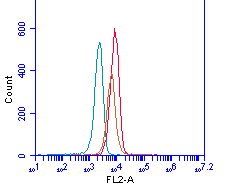 |
| 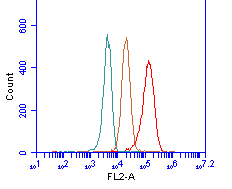 | 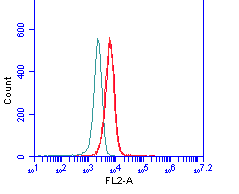 |

**Supplementary Figure 3.** Additional flow cytometry histograms of PPC1 and M21 cells labelled via HRP-mediated proximity biotinylation. Cells were treated and analysed as described in the main figure. (n=3)
